# Kynurenine monooxygenase blockade reduces endometriosis-like lesions, improves visceral hyperalgesia, and rescues mice from a negative behavioural phenotype in experimental endometriosis

**DOI:** 10.1101/2024.05.13.593856

**Authors:** Ben Higgins, Ioannis Simitsidellis, Xiaozhong Zheng, Frances Collins, Natalie Z.M. Homer, Scott G. Denham, Joanna P. Simpson, Mike Millar, Lyndsey Boswell, Hee Y. Lee, Yeon G. Kim, Kyung H. Park, Larry C. Park, Patrick J. Sweeney, Gerard Feraille, Alessandro Taddei, David Chagras, Thierry Alvarez, Scott P. Webster, Andrew Horne, Philippa T.K. Saunders, Damian J. Mole

**Author notes:** Corresponding Author and Lead Contact: Damian J Mole. Indicates senior authors (Andrew Horne, Philippa T.K. Saunders, Damian J. Mole).

## Abstract

Endometriosis is a common and debilitating neuro-inflammatory disorder that is associated with chronic pain. Definitive diagnosis is based on the presence of endometrial-like tissue (lesions) in sites outside the uterus. Kynurenine monooxygenase (KMO) is a mitochondrial enzyme of tryptophan metabolism that regulates inflammation and immunity. Here, we show that KMO is expressed in epithelial cells in human endometriosis tissue lesions and in corresponding lesions in a mouse model of endometriosis. In mice, oral treatment with the potent KMO inhibitor KNS898 induced a biochemical state of KMO blockade with accumulation of kynurenine, diversion to kynurenic acid and ablation of 3-hydroxykynurenine production. In the mouse model of endometriosis, KMO inhibition improved histological outcomes and endometriosis pain-like behaviours, even when KNS898 treatment commenced one week after initiation of lesions. Taken together, these results suggest that KMO blockade is a promising new non-hormonal therapeutic modality for endometriosis.

## INTRODUCTION

Endometriosis is a life-altering condition that affects approximately 10% of females. It is an oestrogen-dependent neuroinflammatory disorder associated with debilitating pelvic pain, excessive fatigue, gastrointestinal and urinary symptoms, and infertility^1^. Worldwide, 200 million prevalent cases are forecast by 2026. Endometriosis is defined by the presence of endometrial-like tissue (‘lesions’) outside the uterus. Physiological hormonal fluctuations in women induce cyclical episodes of cell proliferation, inflammation, injury, and repair within lesions that favour fibroblast to myofibroblast differentiation and fibrosis^1^. We and others have identified metabolic dysfunction in cells associated with development of endometriosis lesions^2^. At present, therapeutic options are largely limited to surgery (that often needs to be repeated) or medical therapies that target hormonal activity with resultant side effects (block conception, menopausal symptoms)^1^. Patient surveys consistently show frustration with the lack of available treatments that can give long term relief from symptoms including pain, low mood and bloating^3^. Analysis of recent clinical trials directed at endometriosis^1^ has highlighted an unmet need for new, non-hormonal approaches to symptom relief. The studies in the current paper have addressed this need by focusing on an enzyme that is known to play a key role in inflammatory processes that are implicated in the aetiology of endometriosis, but which has not previously been investigated as a target.

Our proposed solution to this unmet medical need is by targeting the enzyme kynurenine 3-monooxygenase (KMO). KMO is a critical regulator of inflammation at multiple organ sites that acts by altering metabolic flux through the kynurenine pathway of tryptophan metabolism^4^. KMO is known to be expressed in non-pathological endometrium^5^, but whether KMO is over-expressed in endometriosis lesions and linked to the severity of inflammation remains to be determined. KMO has been identified as a critical step in converting kynurenine to the cytotoxic metabolite, 3-hydroxykynurenine (3HK), that is an oxidative stressor, causes protein cross-linking, and regulates the immune-metabolic interface^4^. Although there is no specific information about a direct role of KMO in endometriosis, there is evidence of dysregulated tryptophan metabolism in a recent study using a preclinical non-human primate model of endometriosis^6^, and increased kynurenine pathway flux at the immune-metabolic interface between stromal cells and NK immune cells in endometriosis lesions^7^.

At present, there is a scientific rationale for KMO inhibition in endometriosis, but it remains to be shown in preclinical experiments whether KMO inhibition is efficacious in decreasing lesion volume or behavioural symptoms which are used as a surrogate for pain responses in model systems. KNS898 is a highly specific small molecule KMO inhibitor with potential for use by women with endometriosis, based on favourable characteristics for oral development in terms of bioavailability and predicted half-life^8,9^. KNS898 is a competitive inhibitor of kynurenine substrate at the active site of KMO with a pIC50 of 8.8^8–10^. We propose that KMO inhibition is a novel therapeutic strategy for endometriosis and, if successful, we will make a significant positive impact for women with this painful, disabling condition. The aim of this project was to obtain proof-of-concept for KMO inhibition as a novel therapy for endometriosis. Specifically, we sought to explore the expression of KMO in biobanked human endometrial and endometriosis lesion tissues, confirm target inhibition of KMO by KNS898 in mice, and define the preclinical efficacy of KNS898 in improving clinical features of disease (specifically hyperalgesia and altered cage behaviour) and reducing endometriosis lesion volume in an experimental mouse model of endometriosis.

## RESULTS

### KMO is expressed in human eutopic endometrium and human endometriosis tissue lesions

To explore whether we could detect variations between expression of KMO in endometrium (eutopic) within the uterus and a variety of lesions obtained from patients, we conducted detailed immunohistochemistry with a primary antibody specific for KMO. On fixed tissue sections of normal human endometrium KMO expression was most striking in epithelial cells lining the glands (Figure 1a, insert B) with lower levels in the luminal layer (insert C). Notably expression in the glands was not uniform (**Figure 1a**). KMO was also strongly immunopositive in human peritoneal endometriosis lesions (**Fig. 1d to Fig. 1g**), and evidently mostly localised to the epithelial tissues surrounding the distended endometrial gland-like structures (DEGLS) (**Fig. 1e and Fig. 1g**). Expression in the stromal compartment appeared variable. In human ovarian endometriosis lesions, KMO was present at low expression levels in the mesothelial layers (**Fig. 1h and Fig. 1i**). Duplex immunohistochemistry with cell phenotype markers CD68 (macrophages) did not show KMO colocalising with these immune cells (data not shown)

**Figure 1.**
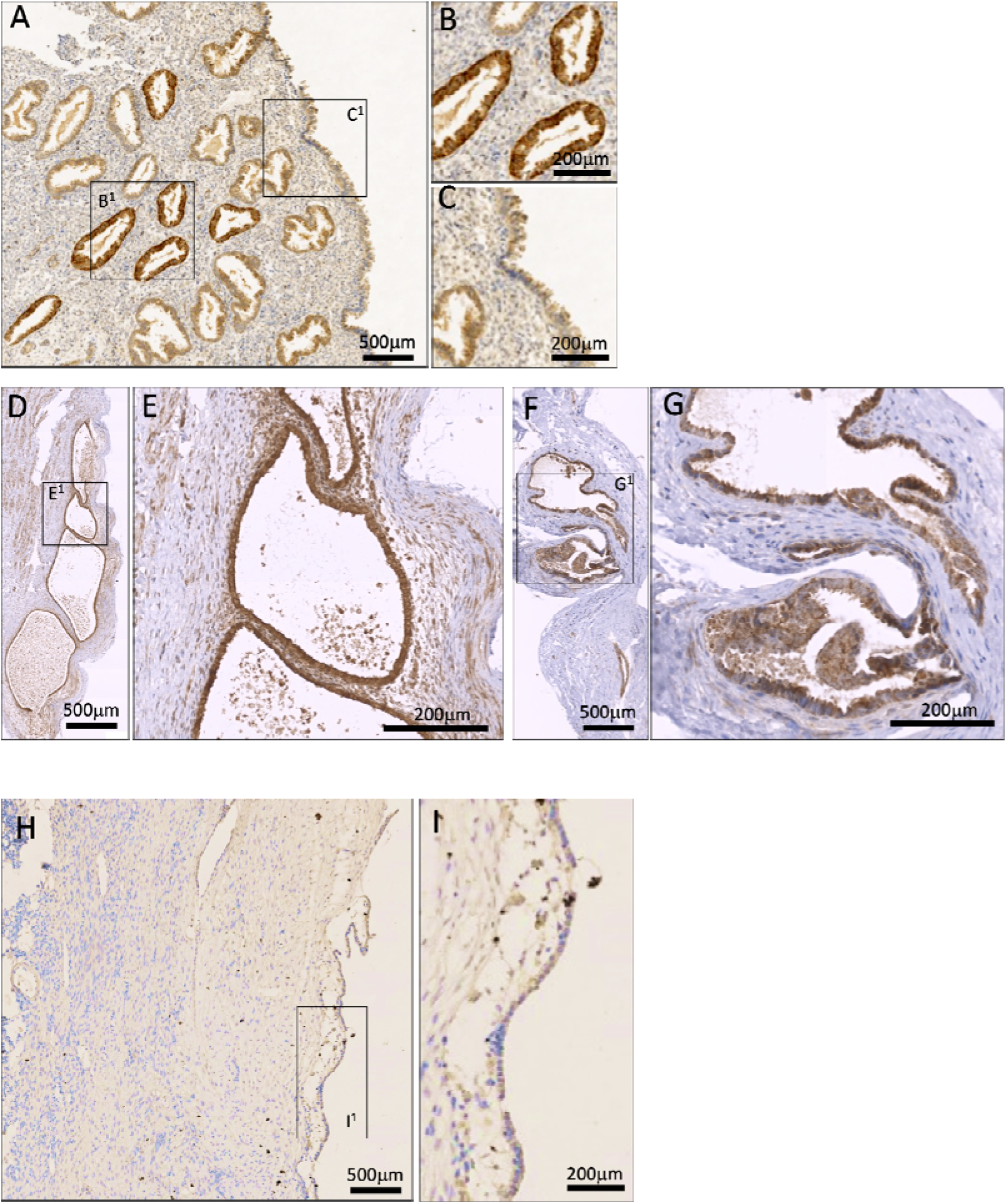
Immunohistochemistry of KMO expression in human endometrium and distended endometriosis gland-like lesions. Fixed tissue sections were stained with anti-KMO antibody (1:500 dilution) and visualized with DAB as described in the Methods section. **Panel A.** Normal human endometrium (200 X magnification); **B^1^** and **C^1^** insets denote areas shown in panels **B** and **C** at higher magnification. KMO expression is demonstrated as dark brown DAB-positive staining and is most intense in the epithelial cells lining the glands. **Panels D through G.** Human peritoneal endometriosis tissue lesions stained with anti-KMO antibody visualized with DAB. D/**E** PIN3652, Stage II, F/**G** PIN3306 Stage I. Note intense staining of cells lining glandular structures. **Panel H.** Ovarian-type endometriosis tissue lesion with higher magnification inset (**I^1^**) showing KMO expression present but at lower intensity in the mesothelial tissue surface (this sample does not have epithelial cells in the lesion).

### Oral KNS898 inhibits KMO in mice

Next, we established that oral dosing of KNS898 by gavage in mice resulted in inhibition of KMO. Using n=3 mice per group, we administered KNS898 at 0.01 mg/kg, 5 mg/kg, and 25 mg/kg twice daily (b.d.) in vehicle for seven days, as described. Plasma drug levels and metabolite concentrations are shown in **Figure 2**. KNS898 dosed at 0.01 mg/kg b.d. resulted in a mean (± S.E.M.) plasma drug level of 0.18 ± 0.01 ng/mL, 5 mg/kg resulted in 88.8 ± 22.6 μg/mL, and 25 mg/kg gave 483.9 ± 84.0 μg/mL. The difference between groups was statistically significant by one-way ANOVA with post hoc Tukey’s test (P = 0.001) (**Fig. 2a**). KMO blockade with KNS898 was clearly measurable. A backlog in the KMO substrate KYN was evident: KNS898 dosed at 0.01 mg/kg b.d. resulted in a mean (± S.E.M.) plasma level of KYN of 339 ± 39 ng/mL, 5 mg/kg resulted in 4940 ± 483 ng/mL, and 25 mg/kg gave 3682 ± 634 ng/mL. The difference between groups was statistically significant by one-way ANOVA with post hoc Tukey’s test (P = 0.001). The increase in KYN at maximal inhibition was approximately 14-fold compared to the level seen after KNS898 0.01 mg/kg (**Fig. 2b**). Excess KYN was metabolised to KA by kynurenine aminotransferase: KNS898 dosed at 0.01 mg/kg b.d. resulted in a mean (± S.E.M.) plasma level of KA of 629 ± 103 ng/mL, 5 mg/kg resulted in 14399 ± 3394 ng/mL, and 25 mg/kg gave 15965 ± 789 ng/mL. The difference between groups was statistically significant by one-way ANOVA with post hoc Tukey’s test (P = 0.001). The fold increase in KA at maximal inhibition was approximately 25-fold compared to the level seen after KNS898 0.01 mg/kg (**Fig. 2c**). KMO blockade resulted in a statistically-significant reduction of 3HK in plasma: KNS898 dosed at 0.01 mg/kg b.d. resulted in a mean (± S.E.M.) plasma level of 3HK of 27.7 ± 7.2 ng/mL, 5 mg/kg resulted in 4.2 ± 0.3 ng/mL, and 25 mg/kg gave 0.9 ± 0.4 ng/mL. The difference between groups was statistically significant by one-way ANOVA with post hoc Tukey’s test (P = 0.001). The fold decrease in 3HK at maximal inhibition was approximately 30-fold compared to the level seen after KNS898 0.01 mg/kg (**Fig. 2d**). Overall, there was a clear dose response to KNS898 administration leading to maximal KMO blockade at 25 mg/kg b.d. This dose was therefore selected for efficacy experiments going forward. A diagrammatic representation of the kynurenine pathway is shown as Figure 2e for reference.

**Figure 2.**
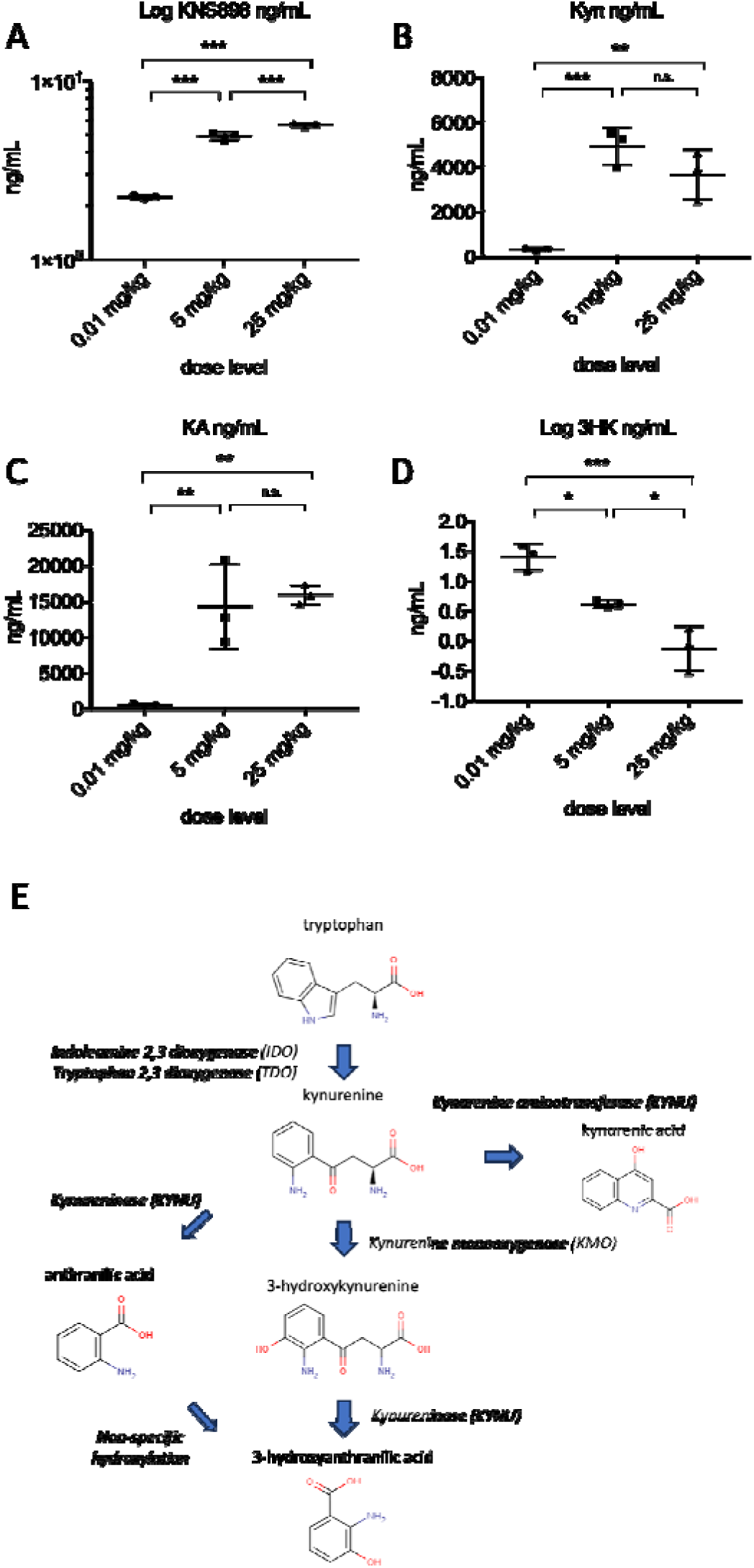
KNS898 plasma levels and pharmacodynamic effect of KMO blockade. Mice (n=3 per group, individual data shown) were given KNS898 twice daily by gavage at the doses shown. After 7 days, blood was sampled at euthanasia and KNS898 levels and kynurenine pathway metabolite levels measured by LC-MS/MS. **A.** KNS898 drug levels. **B.** Kynurenine. **C.** Kynurenic acid. **D**. 3-hydroxykynurenine (logarithmic scale). Comparison between groups by one way ANOVA with post hoc Tukey’s test. *P <0.05, **P<0.01, ***P<0.001, n.s. not statistically significant. **E.** A diagram of the kynurenine pathway showing the key step catalyzed by KMO.

### KMO blockade reduces endometrial gland-like lesion burden in experimental endometriosis in mice

The experimental design for the mouse model of endometriosis is shown in **Figure 3a**. The pharmacological effect of KNS898 therapy showed appropriate levels of KNS898 detected in plasma (**Fig 3b**), with accumulation of kynurenine (**Fig. 3c**), inhibition of 3HK production (**Fig. 3d**) and diverted metabolism of accumulated kynurenine to kynurenic acid (**Fig. 3e**). All recipient mice inoculated with donor tissue (groups G3, G4, and G5) developed distended endometrial gland-like structures (DEGLS). The incidence of DEGLS formation was enumerated at autopsy, and the axial length of each DEGLS was measured after excision from the surrounding tissue. In G3 (endometriosis + vehicle), 8 of 15 (53%) of the inoculated animals had developed DEGLS. In KNS898-treated group G4 (endometriosis + treatment from Day 19), DEGLS formed in 4 of 15 mice (26.7%) and in G5 (Endo + treatment start on Day 26) in 6 of 15 mice (40%). As expected, no DEGLS were formed in the non-inoculated control and sham groups. The total number of DEGLS per animal in each group was highest in G3 with an average of 4.0 per animal with DEGLS (total = 32 DEGLS in 8 mice in G3). Mice with endometriosis receiving KNS898 from the time of inoculation (G4) had an average of 2.0 DEGLS per animal with DEGLS (total = 8 DEGLS in 4 mice in G4) and those receiving KNS898 1 week after inoculation (G5) had an average of 1.8 DEGLS per animal (total = 11 DEGLS in 6 mice in G5) (**Figs. 3f and 3g**). Statistical analysis by ANOVA showed a significant difference in endometriosis DEGLS burden between groups (P = 0.0295 for DEGLS per animal; P = 0.004 for DEGLS per group). DEGLS axial length and derived volume did not differ between groups (**Supplementary Fig. 1a and b**). All recipient mice inoculated with donor tissue lost body weight following inoculation which then gradually recovered. After recovery, body weight of all three inoculated groups was lower compared to the control groups for the duration of the study. Overall, there was no significant difference in body weight between G3 and the KNS898 treatment groups G4 and G5 (endometriosis + treatment from Day 26) (**Suppl. Fig. 1c**).

**Figure 3.**
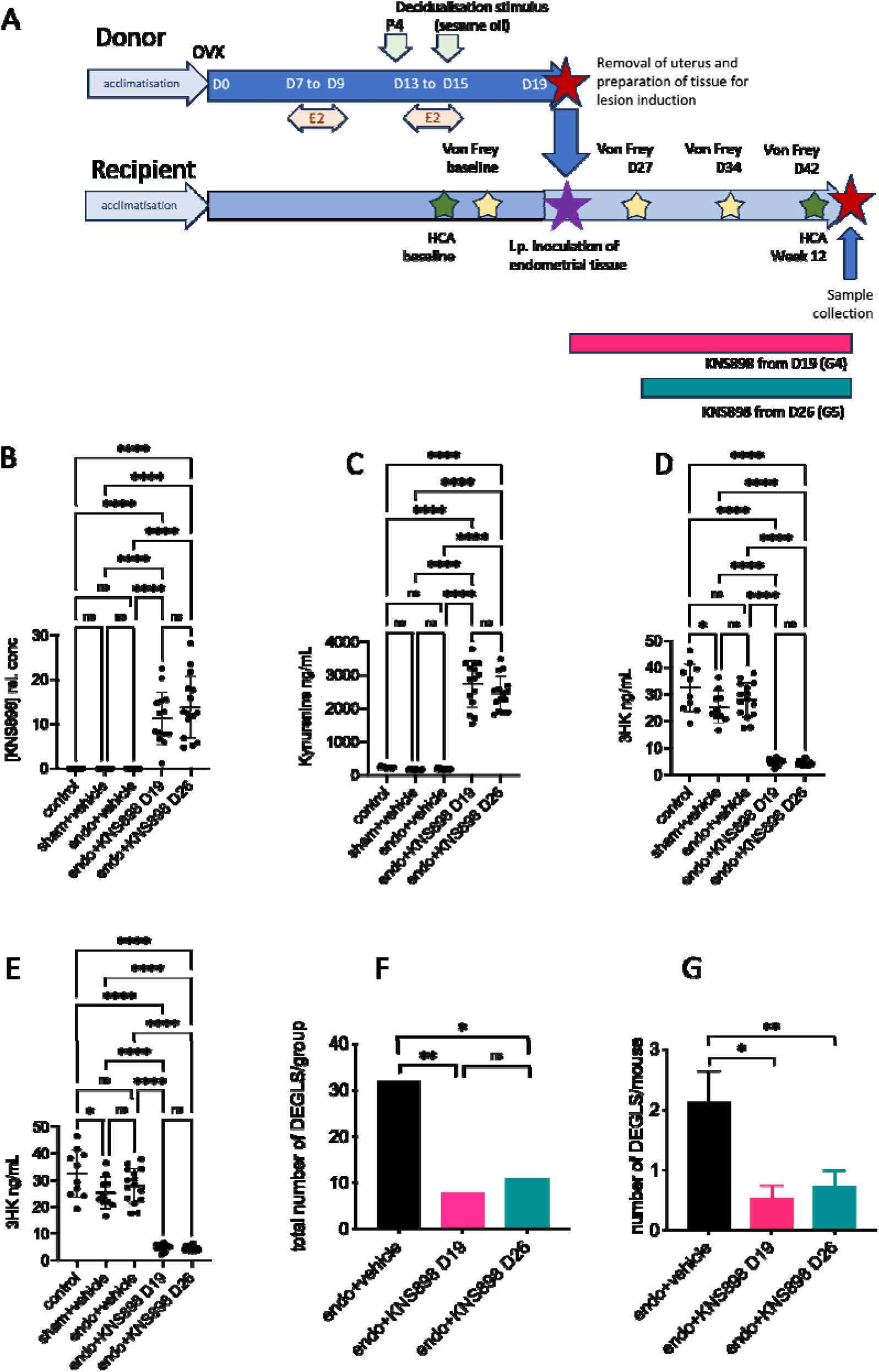
Therapeutic effect of KNS898 in an experimental mouse model of endometriosis. **A.** Experimental design. Ovariectomized (OVX) donor mice were hormonally stimulated as shown (E2 estradiol, P4 progesterone). At Day 19, donor mouse endometrial fragments were inoculated into recipient mice in a 1:1 ratio. KNS898 treatment 25 mg/kg twice daily by oral gavage was commenced at Day 19 or after a 1 week interval on Day 26 and in both cases continued for 2 weeks. Groups were G1: n=10, control mice; G2: n=10, sham-operated control mice; G3: n=15, endometriosis + vehicle; G4: n=15, endometriosis with KNS898 commenced at Day 19; G5: n=15, endometriosis with KNS898 commenced at Day 26. **B.** KNS898 drug levels (relative concentrations). **C.** Kynurenine. **D.** 3-hydroxykynurenine. **E**. Kynurenic acid. Individual data are shown in panels B through E. **Panels F and G.** Enumerated distended endometriosis gland-like structures (DEGLS) in recipient mice by treatment group. **F**. Total number of DEGLS per group. **G**. Total number of DEGLS per animal for all animals in the group; bars show mean with s.e.m. (**G**). Comparison between groups by one way ANOVA with post hoc Tukey’s test. *P <0.05, **P<0.01, ***P<0.001, ****P<0.0001, n.s. not statistically significant.

### KMO is expressed in experimental endometriosis in mice

Histological examination of DEGLS identified them as containing cystic structures lined with epithelial layers identifiable as columnar epithelium, pseudostratified epithelium, squamous epithelium, and cuboidal epithelium, with goblet cells. These DEGLS were considered to represent endometriosis-like lesions derived from the implanted basal endometrial/myoepithelial layers of the donor mice uteri (**Suppl. Fig 2**). Immunohistochemistry using an antibody to KMO showed KMO protein expression localised mainly to the epithelial cells lining of the DEGLS, with a lesser degree of KMO positive staining in the closest surrounding connective tissue, in keeping with the previously observed KMO expression pattern in human endometriosis lesion tissue (**Fig. 4a and 4d and Supplementary Fig S3**). The thickness (area divided by length) of the KMO positive epithelial layer was quantified for each DEGLS section using QuPath and there was no difference between groups G3, G4 and G5 (**Fig. 4b and 4e**). However, quantification of KMO expression confirmed the high intensity of KMO staining in the epithelial lining layers (**Fig. 4c and 4f**), but also showed a clear and statistically significant reduction in KMO expression intensity in those areas in DEGLS removed from mice treated with the KMO inhibitor KNS898 (**Fig. 4g**; P = 0.008). Representative micrographs from each of groups G3, G4 and G5 are presented in **Figures 4h, 4i and 4j**.

**Figure 4.**
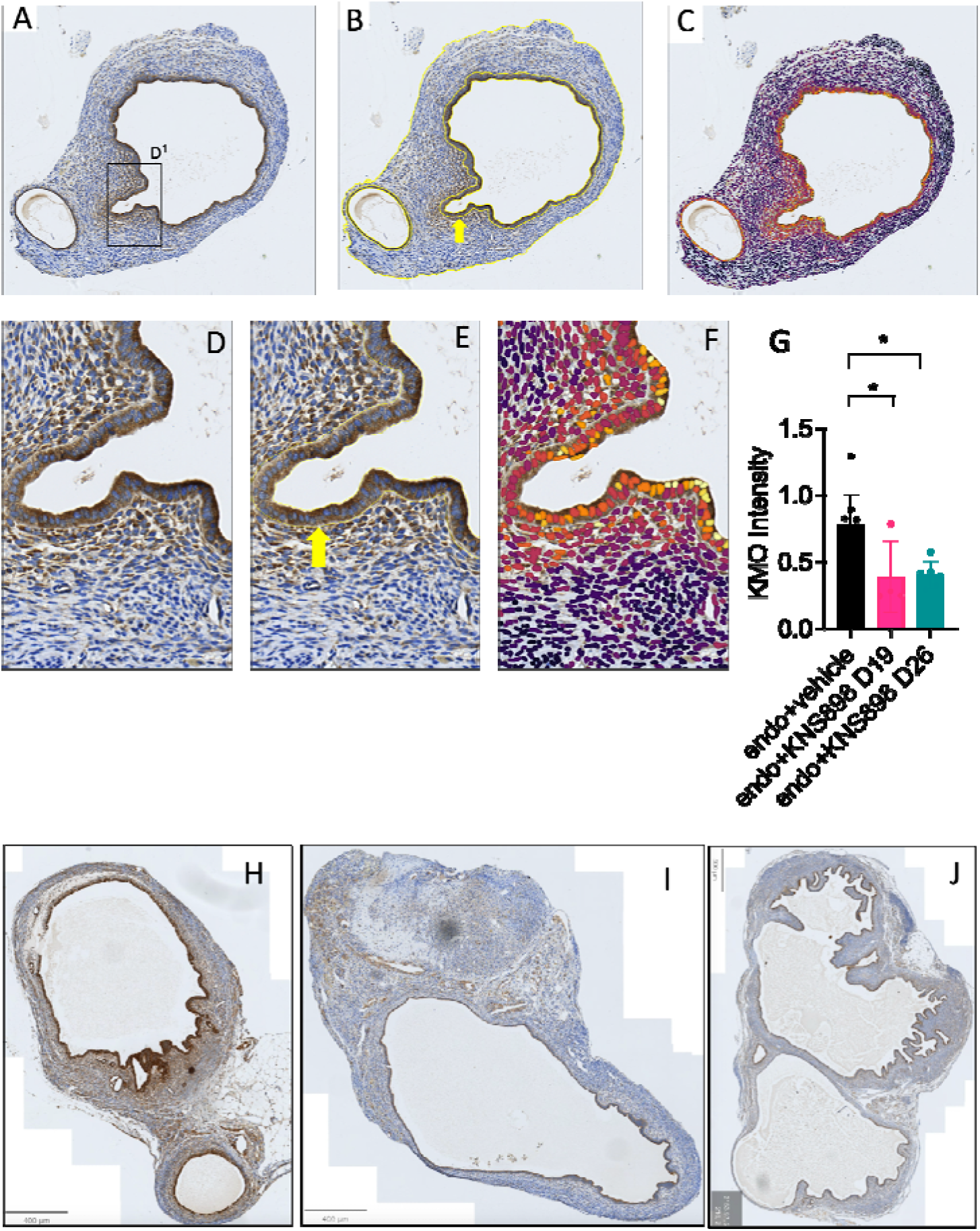
Quantification of KMO expression in mouse model distended endometriosis gland-like structure (DEGLS) lesions. Sections were visualized at 200 X magnification (**A, B, C**) with higher magnification insets shown (**D, E, F**). **Panel A and D.** Fixed tissue sections were stained with anti-KMO antibody (1:500 dilution) and visualized with DAB as described in the Methods section. **B and E.** QuPath was used to identify the epithelial tissue layers (yellow arrows denote the boundary) which were quantified by thickness. **C and F**. KMO expression intensity quantified and heat map expression values are overlayed. **G.** KMO expression staining intensity per unit area of endometriosis DEGLS epithelium, categorized by treatment group (G3 endometriosis + vehicle; G4 endometriosis + KNS898 from D19; G5 endometriosis + KNS898 from D26. Individual data points shown. Comparison between groups by one way ANOVA with post hoc Tukey’s test. *P <0.05.. **H, I and J.** Representative micrographs from G3 (**H**), G4 (**I**) and G5 (**J**) showing KMO expression in the epithelial cells lining each DEGLS.

### KMO inhibition reduces mechanical allodynia in experimental endometriosis

Clinical endometriosis is associated with visceral hyperalgesia and central sensitisation to pain^11^. Visceral and central hyperalgesia may be tested in rodents using the Von Frey filament test^12^. Baseline reaction values for hind paw and bladder Von Frey tests showed no significant difference in mechanical allodynia before inoculation. In established endometriosis without treatment (group G3), the mechanical allodynia threshold in the hind paw was statistically significantly lower compared to baseline for the group. When compared to the control groups at the corresponding time point beginning 1 week after inoculation and continuing until the end of the study. Day 26 KNS898-treated group (G5) showed a statistically-significant improvement in mechanical allodynia in the hind paw using the Von Frey test compared to mice in G3 with untreated endometriosis given vehicle control (Two-way ANOVA, Group effect P = 0.003, time effect P < 0.0001; Dunnett’s multiple comparison test G5 vs G3 P=0.001) (**Fig. 5a**). The mechanical allodynia threshold for the bladder reflex also was lower in mice with endometriosis compared to baseline throughout the study, and KNS898 treatment commencing at D26 (G5) was associated with a statistically significant improvement in bladder mechanical allodynia threshold at D42 compared to mice with untreated endometriosis given vehicle control (G3)(Two-way ANOVA, Group effect P = 0.038, time effect P < 0.001; Dunnett’s multiple comparison test G5 vs G3 P=0.021)(**Fig. 5b**).

**Figure 5.**
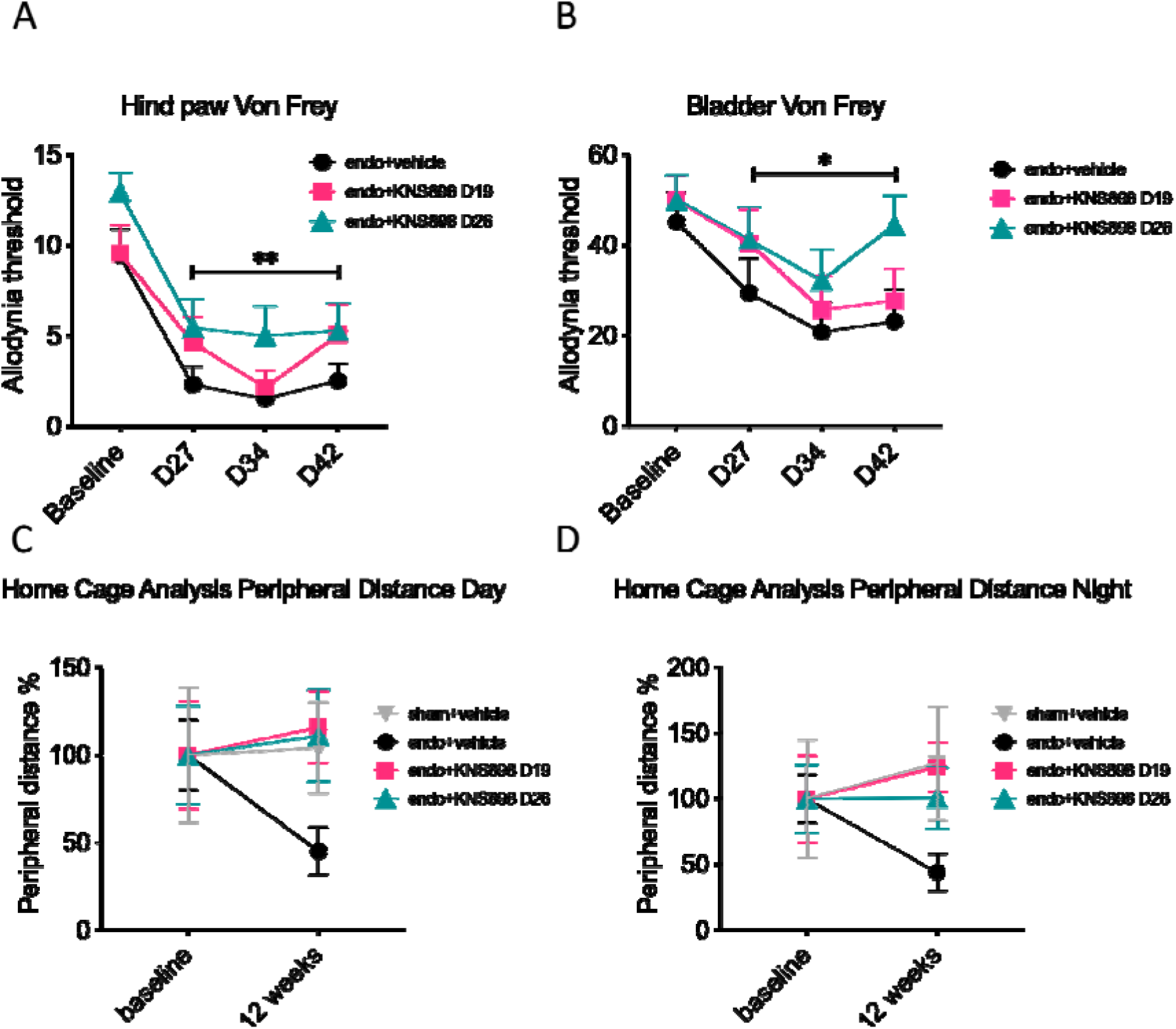
Effect of KNS898 on mechanical allodynia and illness behaviour in an experimental mouse model of endometriosis. **A.** Hind paw Von Frey filament test showing effect of endometriosis and KNS898 treatment in groups G3, G4 and G5 **B.** Bladder Von Frey filament test **C.** Home Cage Analysis of motility showing a daytime motility deficit in mice with endometriosis compared to control mice, and clear restitution of normal motility in KNS898 treated groups. **D**. Nighttime home cage motility analysis showing the benefit of KNS898 treatment on normalizing the motility deficit seen in mice with endometriosis. Data are mean with s.e.m. For A and B, statistical comparison between groups was by two-way ANOVA to compare Group effect and Time effect, with multiple group comparison using Dunnett’s T3 test. Asterisks represent treatment group effect statistical significance of the Dunnett’s T3 test comparing treatment group G5 to G3 vehicle control *P <0.05, **P <0.01. For C and D, Welch’s ANOVA with multiple group comparison using Dunnett’s T3 test was used. Although the ANOVA was statistically significant, the post hoc Dunnett’s T3 was not, therefore no asterisks are marked.

### KMO inhibition rescues impaired cage exploration behaviour and mobility in mice with endometriosis

HCA peripheral moving speed, time at cage edge, and illness behaviour, including temperature, motility and cage exploration behaviour was quantified using Home Cage Analysis (HCA). Baseline HCA was recorded before inoculation and at the end of the experiment. Mice with endometriosis without treatment showed an overall reduction in activity in moving distance and moving speed relative to baseline, and compared to sham-operated control groups, indicating a negative effect on behaviour due to endometriosis. Importantly, mice with endometriosis treated with KNS898 showed marked improvement in motility and cage exploration behaviour compared to untreated endometriosis mice, and although this difference between groups was statistically significant by Welch’s one-way ANOVA, post hoc testing (Dunnet’s T3) was not significant between groups. This qualitative difference of time spent exploring the periphery of the cage being lower for mice with endometriosis treated with vehicle control was seen in both the day and night phases (**Figs. 5c and 5d**). A similar improvement with KNS898 treatment compared to vehicle control mice with endometriosis was seen for total moving distance, total moving speed, and peripheral distance, but not for total moving or climbing time, for which no difference between groups was detected. Together, these data indicate that KMO inhibition with KNS898 results in an improvement in well-being evidenced by improved cage exploration behaviour in addition to improved objective histological measures of endometriosis disease burden.

## DISCUSSION

In this study, we set out to investigate the potential for KMO inhibition as a non-hormonal therapy for endometriosis. First, we confirmed that KMO was expressed in human endometrium by immunohistochemistry, and then showed that KMO was clearly expressed in the epithelial cells in human endometriosis lesions. Next, we demonstrated that the highly specific KMO inhibitor KNS898 was orally bioavailable when given twice daily by gavage in mice, and clearly blocked KMO activity in a dose-dependent manner at a dose of 25mg/kg. We therefore used that dose to test the efficacy of KMO blockade with KNS898 in mice with experimentally-induced endometriosis. One important finding of this project is that KMO blockade resulted in a reduction in endometriosis severity compared to untreated mice with endometriosis, specifically in terms of reducing i) the number of mice that developed endometriosis tissue lesions, and ii) the number of lesions per mouse in those that did develop lesions. The histopathology of the experimental endometriosis lesions was sufficiently similar macroscopically to that seen in the examined human DEGLS, and KMO was evidently highly expressed in the same tissue distribution in model lesions compared to human disease. KMO blockade also decreased lesion KMO expression. Critically, and importantly from a translational perspective, therapeutic blockade of KMO improved visceral hyperalgesia measured by reduced mechanical allodynia and restored normal cage exploration behaviour and mobility in treated mice compared to untreated mice with endometriosis. Together, these data show that KMO is expressed in human and mouse endometriosis tissue lesions and that therapeutic KMO blockade reduces the number of endometriosis lesions and improves holistic metrics of disease behaviour in mice.

The model of endometriosis, using inoculation of endometrial tissue of ovariectomized donor mice, reliably induced the pathophysiological symptoms indicative of endometriosis in recipient mice. Test groups inoculated with endometrial tissue (G3-G5) showed significant growth of ectopic endometrial tissue. Groups treated with test article experienced significantly less DEGLS development (significantly fewer DEGLS were noted in treated groups when compared to vehicle treated groups). Disease burden in the treatment group that had treatment starting immediately after inoculation was lower than that of the vehicle-only treated group. It is not clear why mean cystic size and cystic volume in treated animals was not smaller in treated animals. We can only speculate that KMO blockade may potentiate rapid involution of cysts, but this cannot be proven mechanistically here.

Mice that received inoculated endometrial tissue showed a measurable and increased visceral hyperalgic pain response (lower mechanical threshold) as measured by bladder response to von Frey filament testing, and improvement in a surrogate marker of central sensitisation to pain measured by hind paw Von Frey filament testing when compared to control mice. The mechanical threshold of both treatment groups trended higher compared to the vehicle treated group. One interpretation of these data is that KMO inhibition reduced responses to pain caused by the presence of endometriosis, i.e. improving visceral hyperalgesia.

Home cage behaviour using HCA indicated a reduced overall activity in endo-inoculated mice when compared to control mice. It should also be noted that test article-treated animals in both groups showed more locomotor behaviour and a seemingly better quality of life within the home cage environment when the home-cage dynamics were monitored. This supports the notion that treated mice exhibit less propensity for behaviours that are, at times, typical of depressive and anxiety-like behaviour in home-cage, group housed conditions.

Under the experimental conditions imposed, treatment with KNS898 on two dosing schedules provided a significant reduction in DEGLS formation within the inoculated mice, as well as a seemingly higher pain threshold. This attenuation of pain response was coupled with increased activity levels in the home cage. Taken together, these data suggest a therapeutic effect to alleviation of certain salient symptoms of endometriosis, as well as a reduction in the number and size of DEGLS.

Non-pathological endometrium is a site of high KMO expression. Because endometriosis lesions in women are ‘endometrial-like’ tissue rather than normal endometrium, we tested, and demonstrated expression of KMO in the epithelial layers of endometriosis lesions sampled from women undergoing surgery for endometriosis. First-line medical treatment for endometriosis is the contraceptive pill or other ovarian steroid hormone suppressive drugs. Treatment failures are frequent, side effects are common, all approaches are contraceptive. Many women opt for invasive surgery to remove or ablate the endometriosis lesions.

In conclusion, KMO is expressed in human endometriosis tissue lesions and in a mouse model of endometriosis in the epithelial layers of distended endometrial gland-like structures. Oral KNS898 reliably induced a biochemical state of KMO blockade with accumulation of kynurenine, diversion to kynurenic acid and ablation of 3-hydroxykynurenine production. KMO blockade improved histological and symptomatic behavioural endometriosis disease features with an overall benefit, even when treatment commenced one week after establishment of the disease. KMO blockade is therefore a promising avenue for a new non-hormonal therapeutic modality for endometriosis.

## MATERIALS AND METHODS

### Ethical approvals and permissions

The human tissue samples were obtained from participants who had given fully informed written consent under ethical approval granted by Lothian Research Ethics Committee (LREC 11/AL/0376). Human tissue samples were obtained with ethical approval and fully informed consent from individuals attending the Royal Infirmary of Edinburgh as described below. Animal experiments conducted by NAASON Inc were carried out according to the National Institute of Health (NIH) & National Institutes of Health Korea (NIHK) guidelines for the care and use of laboratory animals and approved by Naason Science in accordance with all applicable FELASA, IACUC and AAALAC guidelines. Animal experiments outsourced to Syneos Health were conducted with institutional ethical approval.

### Human Patients and Samples

Tissue samples were collected from patients undergoing a diagnostic laparoscopy for suspected endometriosis following Endometriosis Phenome and Biobanking Harmonisation Project (EPHect) guidelines^13^. Patient summary characteristics are presented in **Supplementary Table S1**. Note there was a range of disease stages assigned at time of surgery according to American Fertility Society (AFS) criteria^14^. Cycle stage was determined by measuring hormones in blood according to standard protocols and assessment of eutopic endometrial tissue histology when such samples were available^15^. Lesions were recovered from 17 patients, of these n=10 were recovered from the peritoneal side wall consistent with classification as superficial peritoneal endometriosis lesions and n=5 from the cysts of ovarian disease (endometrioma) (n=2 on hormones). Eutopic endometrium was from 4 patients n=3 of which had no lesions at time of surgery (noted as stage 0). General histology of samples was assessed using H&E staining.

### Immunohistochemistry (human endometrium and human endometriosis tissue lesions)

5 μm sections of formalin-fixed paraffin-embedded tissue blocks were mounted on SuperFrost Plus adhesion slides (Thermo Fisher Scientific). Sections were deparaffined with xylene and rehydrated prior to heat-induced antigen retrieval using Instant Pot: Tris-EDTA pH9^16^. Sections were washed with tap water and incubated in phosphate buffered saline (PBS) for 5 minutes. Endogenous peroxidase was blocked with 0.3% hydrogen peroxide in 70% v/v methanol for 30 mins at room temperature then washed in PBS prior to blocking in Normal Goat Serum (NGS)/PBS/bovine serum albumin (BSA)(5%) for 30 mins and streptavidin for 15 mins. Sections were washed twice in PBS and then blocked with biotin for 15 mins and washed in PBS. The primary antibody to KMO (KMO Rabbit polyclonal, Proteintech, Catalog Number:10698-1-AP)^17^ was diluted to a final concentration of 1:1000 in NGS/PBS/BSA and incubated overnight at 4°C in a humidity chamber. Sections were washed twice with PBS/Tween 0.05% (1ml Tween in 2L PBS) for 5 mins. The secondary detection antibody Goat Anti-Rabbit Biotinylated (Vector Cat number: BA-1000) was diluted in NGS/PBS/BSA (1:500) and incubated for 30 mins, prior to washing twice in PBS/Tween 0.05%, for 5 mins before adding the detection system reagent Streptavidin-HRP (DAKO Cat Number P0397) 1:500 in PBS for 30 min, washed and stained with DAB (DAKO Cat Number K3468) as per manufacturer’s directions and incubated for 5 mins before a final wash with tap water. Sections were counterstained with Haematoxylin, dehydrated through graded ethanol and mounted. Sections of stained slides were scanned on a Zeiss Axioscan Z1 slide scanner and exported as TIFF files: images were evaluated for stromal, epithelial and immune cell content.

### KNS898 preparation for oral administration

KNS898 powder was weighed and dissolved at the required concentrations in a final vehicle of 2% DMSO, 20% PEG200, 78% 0.15M NaCl by volume. Brief sonication on ice was done to facilitate disolution.

### In vivo confirmation of KMO inhibition by KNS898 in mice

This experiment was outsourced to Syneos Health (Les Templiers, 2400 route des Colles, 06410 Biot, Sophia-Antipolis, France). A formal pharmacokinetic/pharmacodynamic study was not required at this stage. Female C5Bl/6J mice aged 12 weeks were purchased from Charles River Laboratories, maintained on standard 12 hour light-dark cycle, given free access to water and standard chow before being randomised to one of three dose levels of KNS898 (n=3 per group, total n=9 mice). Dose levels tested were 0.01mg/kg, 5mg/kg and 25mg/kg. Mice were gavaged with 0.5 mL of drug in vehicle twice daily for 7 days before euthanasia and plasma sampling.

### Plasma samples

Blood was sampled into Sarstedt Microvette CB K2EDTA 300 μL tubes and centrifuged at 5,000 rpm (2380 RCF) for 3 mins. Plasma was aliquoted, frozen on dry ice and transferred to storage at −80°C prior to temperature-controlled shipping.

### LC-MS/MS analysis of plasma drug levels and kynurenine metabolites

Plasma samples (100 μL) were diluted at a 1:1 ratio with 4% phosphoric acid and enriched with 50 ng ^13^C_6_-kynurenine, ^13^C_6_-3-hydroxykynurenine (Sigma Aldrich, custom synthesis) and d5-kynurenic acid (CDN isotopes). 12-point calibration standards (0.1 to 100 ng) were prepared for KNS898, kynurenine (KYN), kynurenic acid (KA), and 3-hydroxykynurenine (3HK) and extracted alongside samples using solid phase extraction plates (Waters Oasis HLB, 10 mg sorbent, 30 μm particle size). Extracts were dried down under nitrogen and reconstituted in LC-MS grade water (100 μL). 10 μL was injected onto a column (Ace C18-PFP column; 100 x 2.1 mm internal diameter 1.7 μm; HiChrom (VWR, Lutterworth)) using an Acquity I-Class UPLC liquid chromatography system (Waters) linked to a QTRAP 6500+ mass spectrometer (AB Sciex)^10^. The flow rate was set at 0.4 mL/min with a column temperature of 40°C. Separation was carried out using a gradient mobile phase system of A – 0.1% aqueous formic acid and B – 0.1% formic acid in methanol, starting at 15%B, rising to 85%B over 6 mins and returning to 15%B by 9 mins. Mass spectrometry settings were for positive mode electrospray (5.5 kV, 700°C) and multiple reaction monitoring *m/z* 209.0 → 192.2 for KYN, *m/z* 189.9 → 144.1 for KA *m/z* 225.0 → 208.0 for 3HK and *m/z* 361.1 → 120.1 for KNS898 and for internal standards were *m/z* 231.0 → 214.0 for ^13^C_6_-3HK, *m/z* 195.1 → 177.2 for d5-KA and *m/z* 215.0 → 197.8 for ^13^C_6_-kynurenine. Retention times for KYN, KA, 3HK and KNS898 were 1.8, 3.2, 1.2 and 6.5 mins, respectively and 3.2 mins for d5KA, 1.8 mins for ^13^C_6_-3HK and 1.2 mins for ^13^C_6_-KYN. Data were acquired by Analyst 1.7.1 software (AB Sciex) and linear regression analysis was carried out on MultiQuant 3.0.3 software (AB Sciex) where peak integrations and amounts of each kynurenine metabolite and KNS898 were calculated using the peak area ratio of compound/internal standard, with data further handled in Microsoft Excel 2016 as described^18^.

### Experimental mouse model of endometriosis

This experiment was outsourced to Naason Science Inc., Osong, Korea (KBIO New Drug Development Center #506, Chungbuk, Korea, 28160) using protocols originally developed by the Saunders team in Edinburgh^19,20^. Experimental design and groups are shown in **Figure 3a** and **Supplementary Table S2**. There were 5 groups of mice with n=10-15/group):25 mg/kg KNS898 was administered twice a day via oral gavage in two of the groups of mice. Group 4 received KNS898 from the time of endometrial tissue inoculation (Day 19; G4); group 5 commenced dosing 1 week after inoculation (Day 26; G5). Group 3 received vehicle (2% DMSO, 20% PEG200, 78% 0.15M NaCl) in the same regimen.

To perform the mouse model of endometriosis, donor female C57Bl/6 mice aged 6 weeks were acclimatized for 2 weeks prior to surgery. Ovariectomy (Day 0) was performed at 8 weeks of age under general anaesthesia with monitoring, with analgesia that extended to the post-operative period with buprenorphine (0.03 ml) (Vetergesic® 3 mg/ml, Ceva Inc., Korea) subcutaneously. To prepare donor tissue that would best replicate menstrual-like tissue in women, ovariectomized (OVX) mice were primed with daily s.c. injections of 100 ng 17β-estradiol (E2) on days 7, 8 and 9. On days 13 – 19 a silastic progesterone (P4) pellet was implanted subcutaneously. These animals were injected once daily with E2 (5 ng in sesame oil) on days 13, 14 and 15. Decidualization was induced in one uterine horn with an injection of 20 µl sesame oil 4 hours after the last E2 injection. On day 19 (4 days after induction of decidual response), donor mice were killed 4 hours after removal of the P4 pellet. Endometrial tissue was then scraped from the myometrial layer of the decidualized uterine horn, suspended in 500 μl PBS and injected via a XG needle sprayed into the lower abdominal cavity of the recipient mouse under general anaesthesia with monitoring and post-operative analgesia as described. The ratio of donor to recipient mouse was 1:1 (from one donor to one recipient). Recipient mice had intact ovaries to ensure ongoing hormonal stimulation of the injected tissue: group allocations are shown in **Supplementary Table S2**.

### Mechanical allodynia test by the Von Frey method

Abdominal and hind paw Von Frey tests were performed in the recipient animals before inoculation (baseline), and 1, 2, and 3 weeks after inoculation. Mechanical threshold was measured using Von Frey filaments. For the hind-paw, 15, 8, 6, 4, 2, 1.4, 0.6, 0.4 g filaments were used, and for the bladder reflex to filament application to the lower abdomen, 60, 26, 10, 8, 6, 4, 2, 1 g filaments were used. The experimenter was blind to the group allocation in order to reduce bias.

### Cage exploration and behavioural assay using Home Cage Analysis

Home Cage Analysis (HCA) was performed in the recipient animals before inoculation (baseline), and at the late stage of treatment. Recipient mice were randomly housed using a Monte Carlo randomization. All animals had a micro-chip (BioMark, USA) inserted to the abdomen prior to being placed in the home-cage. This procedure does not cause undue discomfort or hamper, in any way, animal movement. Total moving distance, total moving time, moving speed, isolation/separation distance, isolated time, peripheral time, peripheral distance, in centre zones time, in centre zones distance, climbing time, and body temperature were tracked automatically by an ActualHCA™ Home Cage Analyzer (ActualAnalytics Ltd., Edinburgh, UK) and processed with proprietary machine learning and artificial intelligence algorithms.

### Endpoint tissue and plasma sampling

On experimental Day 40, all recipient mice were euthanized, and blood collected via cardiac puncture. Whole blood was collected into heparinised tubes, and plasma was separated by centrifugation (3000 rpm for 15 min) at 4°C. Separated plasma was collected in Eppendorf microtubes, frozen on dry ice and stored at −80°C. Photographs of the abdominal cavity were obtained. Lesions from the abdominal cavity were harvested. DEGLS were dissected from the surrounding abdominal tissue and measured for size and volume. DEGLS volume was measured using the following formula^21^: Volume = long diameter × (short diameter/2)2 × π. DEGLS were fixed in a 4% paraformaldehyde solution and prepared for standard H&E. If more than one DEGLS was present in an animal, DEGLS that were not used for H&E were snap-frozen in liquid nitrogen and stored at −80oC.

### Endometriosis lesion histology

Endometriosis lesion tissue blocks were sectioned at a uniform thickness of 5 μm and were mounted onto a microscope slide. The slide then underwent deparaffination and hydration. Paraffin was removed from the slide using xylene, then hydration through graded ethanol and washing were performed. Slides were then stained with Harris haematoxylin and alcoholic eosin Y and mounted after dehydration and clearing with xylene. The H&E-stained images were visualized using a Slide Scanner (Panoramic scan, 3D HISTECH).

### Immunohistochemistry (mouse DEGLS)

Immunohistochemistry to detect KMO in mouse DEGLS tissue was performed on a Leica Bond III automated immunostaining robot. 5 μm thick sections obtained from FFPE (formalin Fixed Paraffin Embedded) samples mounted on superfrost plus slides were stained as follows. Heat induced epitope retrieval (HIER) was performed using Epitope Retrieval Solution 1 (Leica, ER1 pH 6.0 citrate based solution) for 20 minutes at 99^0^C. Tissue sections were then incubated for 10 minutes in hydrogen peroxide to block endogenous hydrogen peroxidase activity followed by 10 minute blocking with normal goat serum. The primary antibody against KMO (Proteintech 10698-1-AP @1:1000 Rabbit)^17^ was incubated for 1 hour, then incubated with a goat anti-rabbit peroxidase conjugated secondary antibody for 30 minutes prior to visualisation with diaminobenzoate (DAB) using standard protocols.

### Digital slide scanning

Whole sections were scanned using a Zeiss Axioscan Z1 whole slide scanner. The image files (.czi) were batch converted to Tif format for image analysis and quantification, acquisition and batch export used Carl Zeiss Zen v2.5 software.

### Quantitative image analysis

Image analysis was carried out using QuPath v0.4.2. Analysis was standardised by using multiple regions of interest from individual slides and collating them into a training image. To improve stain contrast, overlapping DAB and haematoxylin staining was deconvoluted by manual optimisation of the stain vectors. Individual slide epithelia were annotated, with relative DAB optical densities and annotation shape measurements taken. Epithelial thickness and area were calculated to enable correlation with KMO intensity. To generate the heat map of cell KMO expression, a cell detection was carried out on haematoxylin staining using standard parameters, a 5-100um2 area range, cell expansion of 1um and a threshold of 0.14.

### Statistical analysis

Power calculations were performed using G*Power (v3.1.9.4) software. Input parameters were used: 2-tailed, α-error probability = 0.05, and power (1-β error probability) = 0.80. Continuous variable data were tested for Normality of distribution with a one sample Kolmogorov-Smirnov test. Normally distributed data were analysed by one-way ANOVA with post-hoc Dunnett’s T3 for multiple groups. Data comparing treatment group effects at multiple time-points were analysed by two-way ANOVA with multiple comparison testing by Dunnett’s method. Data comparing multiple groups with unequal variances were analyzed with Welch’s ANOVA. Data not following the Normal distribution were analysed with non-parametric Kruskal-Wallis test. Categorical and proportions data were analysed by Fisher’s exact test. Data were visualised with GraphPad Prism.

## LIST OF SUPPLEMENTARY MATERIALS

Fig. S1 and Fig. S2.

Table S1 and Table S2.

## ACKNOWLEDGEMENTS

We would like to thank the University of Edinburgh MRC Confidence in Concept award team: Andrew McBride, Lorraine Jackson. We thank Susan Bodie, Dave Pritchard from Edinburgh Innovations. We thank all staff and support team members at Syneos Health and NAASON Science Inc.

## AUTHOR CONTRIBUTIONS

Conceptualization: DJM, SPW, PTKS, AH

Methodology: BH, IS, NZMH, SGD, JPS, MM, LB, LCP, PJS, AT, SPW, AH, PTKS, DJM

Investigation: BH, IS, XZ, FC, SGP, JPS, LB, HYL, YGK, KHP, LCP GF, AT, DC, TA

Visualization: BH, IS, KHP, LCP, DJM

Funding acquisition: DJM, SPW, PTKS, AH.

Project administration: DJM, XZ, LCP, PES, AT, PTKS, AH, NZMH

Supervision: MM, LCP, AT, PTKS, DJM

Writing – original draft: DJM, LCP

Writing – review & editing: BH, IS, XZ, FC, NZMH, SGD, JPS, MM, LB, LCP, PJS, AT, SPW, AH, PTKS, DJM.

## DATA AND MATERIALS AVAILABILITY

All data are available in the main text or the supplementary materials.

KNS898 availability is restricted under a Material Transfer Agreement. Please contact the corresponding author in the first instance.

