## Supplemental Table 1 for "Kynurenine monooxygenase blockade reduces endometriosis-like lesions, improves visceral hyperalgesia, and rescues mice from a negative behavioural phenotype in experimental endometriosis"

**SUPPLEMENTARY TABLES**

**Table S1.** Patient characteristics and demographics

| Subject PIN | Block number | AFS Stage of Disease | Tissue type | Age | Cycle Stage | On Hormones | BMI |
| --- | --- | --- | --- | --- | --- | --- | --- |
| 3306 | 2014-2167 | I | Peritoneal lesion | 36 | Mid Secretory | No | 26 |
| 3398 | 2015-0882 | I | Peritoneal lesion | 23 | Proliferative | No | 23 |
| 3405 | 2015-1970 | I | Peritoneal lesion | 24 | Early secretory | No | 24 |
| 3412 | 2015-2935 | II | Peritoneal lesion | 26 | Proliferative | No | 25 |
| 3459 | 2015-3955 | I | Peritoneal lesion | 45 | Proliferative | No | 27 |
| 3495 | 2016-2042 | II | Peritoneal lesion | 36 | Proliferative | No | 20 |
| 3597 | 2017-1394 | I | Peritoneal lesion | 22 | Proliferative | No | 23 |
| 3612 | 2017-2835 | III | Endometrioma | 22 | Proliferative | No | 25 |
| 3631 | 2017-3932 | II | Peritoneal lesion | 36 | Proliferative | No | 25 |
| 3640 | 2017-4560 | II | Peritoneal lesion | 32 | Early Secretory | No | 21 |
| 3652 | 2017-5036 | II | Peritoneal lesion | 31 | Early Secretory | No | 24 |
| 3722 | 2020-0319 | 0 | Eutopic endometrium | 34 | Mid Secretary | No | 35 |
| 3776 | 2020-0133 | 0 | Eutopic endometrium | 22 | Proliferative | No | 29 |
| 3777 | 2020-0135 | IV | Endometrioma | 33 | Progesterone effect | Yes | 28 |
| 3777 | 2020-0134 | IV | Endometrioma | 33 | Progesterone effect | Yes | 28 |
| 3778 | 2019-2818 | III | Endometrioma | 32 | Proliferative | No | 21 |
| 3786 | 2020-0284 | IV | Endometrioma | 32 | Proliferative | No | 32 |
| 3787 | 2020-0138 | I | Eutopic endometrium | 30 | Mid Secretary | No | 22 |
| 3794 | 2020-0278 | 0 | Eutopic endometrium | 32 | Proliferative | No | 33 |
| 3799 | 2020-0661 | III | Endometrioma | 29 | N/A | No | 21 |
